# Monitoring Non-Specific Adsorption at Solid-Liquid Interfaces by Supercritical Angle Fluorescence Microscopy

**DOI:** 10.1101/2022.07.19.500728

**Authors:** Aaron Au, Man Ho, Aaron R. Wheeler, Christopher M. Yip

## Abstract

Supercritical angle fluorescence (SAF) microscopy is a novel imaging tool based on the use of distance-dependent fluorophore emission patterns to provide accurate locations of fluorophores relative to a surface. This technique has been used extensively to construct accurate cellular images and to detect surface phenomena in a static environment. However, the capability of SAF microscopy in monitoring dynamic surface phenomena and changes in millisecond intervals is underexplored. Here we report on a hardware add-on for a conventional inverted microscope coupled with a post-processing Python module that extends the capability of SAF microscopy to monitor dynamic surface phenomena thereby greatly expanding the range of potential applications of this tool. We first assessed the performance of the system by probing the specific binding of biotin-fluorescein conjugates to a neutravidin-coated cover glass in the presence of non-binding fluorescein. The SAF emission was observed to increase with the quantity of bound fluorophore on the cover glass. However, high concentration of unbound fluorophore also contributed to overall SAF emission, leading to over-estimation in surface-bound fluorescence. To expand the applications of SAF in monitoring surface phenomena, we monitored the non-specific surface adsorption of BSA and non-ionic surfactants on a Teflon-AF surface. Solution mixtures of BSA and nine Pluronic/Tetronic surfactants were exposed to a Teflon-AF surface. No significant BSA adsorption was observed in all BSA-surfactant solution mixture with negligible SAF intensity. Finally, we monitored the adsorption dynamics of BSA onto the Teflon-AF surface and observed rapid BSA adsorption on Teflon-AF surface within 10 seconds of addition. The adsorption rate constant (k_a_) and half-life of BSA adsorption on Teflon-AF were determined to be 0.2093±0.002 s^−1^ and 3.312±0.032 s respectively using a pseudo-first-order adsorption equation.

## I. INTRODUCTION

Supercritical angle fluorescence (SAF) microscopy is an innovative imaging tool making in-roads in the fields of cell biology, disease diagnostics, and surface characterization. In SAF, the unique distance-dependent fluorophore emission pattern found on the back focal plane of infinity-corrected objectives can be exploited to reduce far-field fluorescence in total internal reflection fluorescence (TIRF) images ^1, 2^ and reveal the precise axial location of single fluorophores in single molecule localization microscopy ^3, 4^. SAF microscopy has been applied to biological studies of cell membranes ^5^ and bacterial DNA amplification on a micro-lens ^6, 7^. SAF enables the measurement of refractive index of a liquid by exploiting the dependence of critical angle on refractive index, ^8, 9^ and has been theorized to detect the emission polarity of fluorophore and to measure the thickness of thin films^10^. SAF has also been used, albeit rarely, to explore time-dependent phenomena. For example, dynamic SAF measurements were used to characterize amyloid-β peptide-bilayer interactions, revealing the dynamics of bilayer disruption ^11^.

In SAF, fluorophores far from the liquid-solid interface are observed in a small cone of light by a high NA objective that only illuminates a region of the back focal plane (BFP) called the subcritical region, while fluorophores near the liquid-solid interface emit light into the super-critical region of the BFP (Figure 1). Comparing relative intensities across these regions of the BFP allows one to infer the dynamics of fluorophores as they approach the liquid-solid interface.

**Figure 1.**
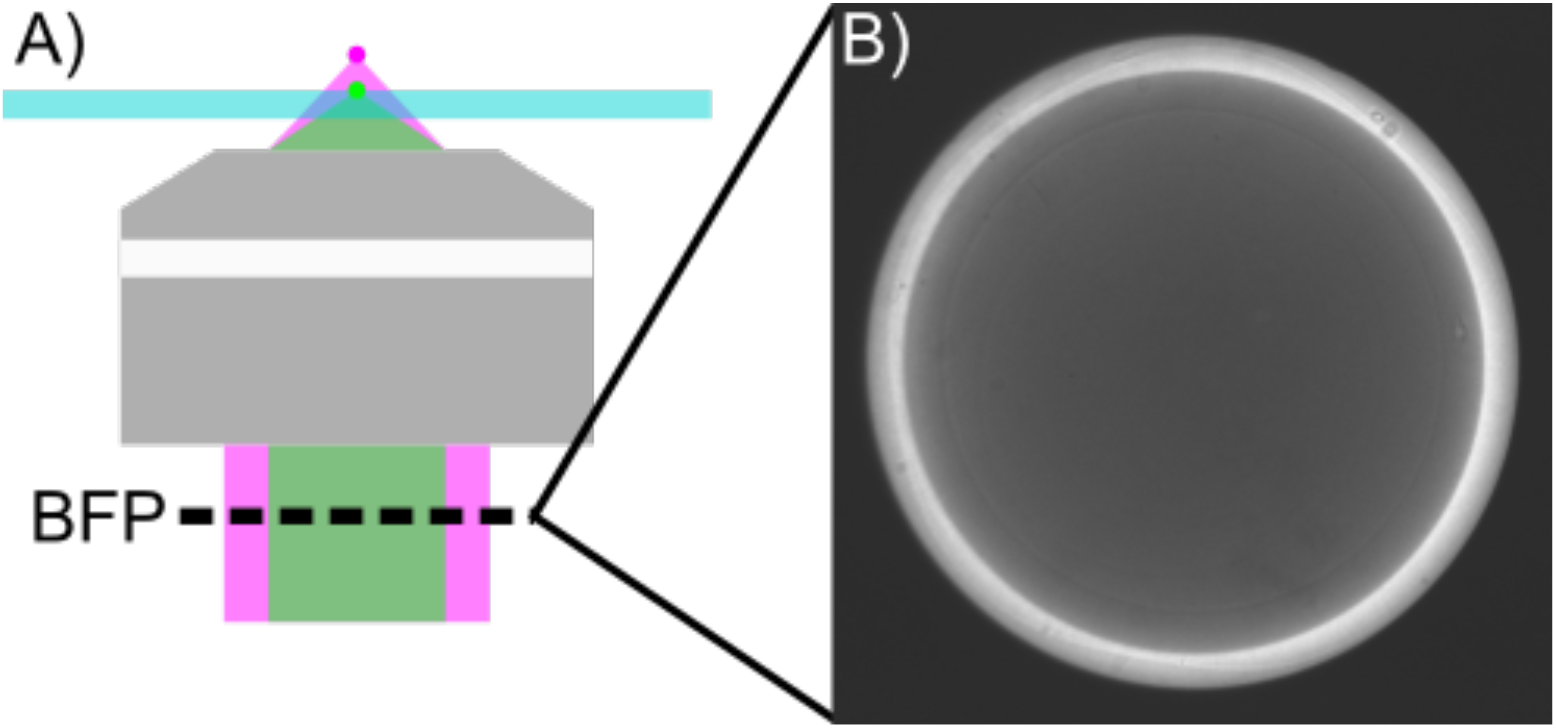
Schematic of Supercritical Angle Fluorescence (SAF) experiment. A) A high NA objective is used to acquire emission fluorescence. Fluorophores located far from the liquid-solid interface produce an illumination pattern within the subcritical region (magenta), while fluorophores near the liquid-solid interface surface (green) create ring-like emission patterns outside of the super-critical radius. B) Representative emission pattern of fluorophores (AF488-BSA) adsorbed onto a glass cover glass.

We report here on an approach for improving the time resolution in SAF microscopy by directly observing the back focal plane using a scientific camera coupled with a dedicated SAF post-processing module to observe interactions at the liquid-solid interface with sub-second time resolutions. Although designed for an Olympus IX83 dual-deck inverted microscope, the SAF hardware module could be readily implemented in other systems. To facilitate the extraction of quantitative SAF measurements, a Python-based post-processing pipeline was developed. In this pipeline, the algorithm detects the circular SAF ring and extracts the SAF intensity profile from either a single image or a stack of images. Additional analysis is then performed to determine the critical angle and differentiate between sub-and super-critical fluorescence. We assessed the performance of this SAF set-up by monitoring the binding of biotin-fluorescein conjugates to a neutravidin-coated surface. To further demonstrate the capability of this platform, we also applied real-time SAF to characterize the dynamics of non-specific adsorption of proteins and surfactants onto Teflon®-AF coated-glass. This work portends the capability of realtime SAF to characterize other near-surface behaviors and dynamics.

## II. METHODS AND MATERIALS

### A. Reagents and Materials

Unless specified otherwise, reagents were purchased from Sigma-Aldrich (Oakville, Canada). Alexa Fluor® 488 conjugated BSA (AF-488 BSA) and Pierce™ biotinfluorescein conjugate were purchased from Thermofisher (Mississauga, Canada). No.1 22×22 mm micro-cover glasses were obtained from VWR International (Mississauga, Canada). Teflon®-AF 1600 polymers were purchased from DuPont (Wilmington, USA). 3D-functionalized neutravidin coverslips, lot # 21-11-10 and Ref # 10403206, were obtained from PolyAn (Berlin, Germany). FC-40 and PFC110 fluoro-solvent were purchased from Sigma-Aldrich (Oakville, Canada) and Cytonix (Bellville, USA). All Pluronic and Tetronic surfactants (BASF Corp., Germany) were generously donated by BASF Corporation (Wyandotte, USA). All experiments using Alexa-488 conjugated BSA were conducted using 76 μM stock solutions of Alexa-488 in Dulbecco’s phosphate buffer saline (DPBS). A 1000 μM stock fluorescein solution was prepared by dissolving appropriated amount of fluorescein disodium salt in DPBS. To prepare 10 μM biotin-fluorescein conjugate stock solution, 5 mg of the biotin-fluorescein conjugate was dissolved in 1 mL of dimethyl sulfoxide and diluted in an appropriate amount of DPBS. All stock solutions of Pluronic and Tetronic were prepared as a 10 % w/w DPBS solutions.

### B. Supercritical Angle Fluorescence Microscope Configuration

For SAF imaging, an optical module was integrated into an Olympus IX-83 dual-deck inverted microscope (Figure 2) [https://github.com/YipLab/IX83-Modules/tree/master/SAF]. In order to ensure even illumination around the BFP and reduce fluorescence contribution from fluorophores far from the glass surface, a ring TIRF configuration was employed. To accomplish this, a custom annulus (7.52 mm inner diameter and 10 mm outer diameter) was placed at the working distance of the first lens of a lens pair. This lens pair (AC254-350-A-ML AC508-200-A-ML, Thorlabs, USA) was used to project the annulus onto the back focal plane of a TIRF objective (UAPON100XOTIRF, Olympus, Japan) generating omnidirectional TIRF illumination.

**Figure 2.**
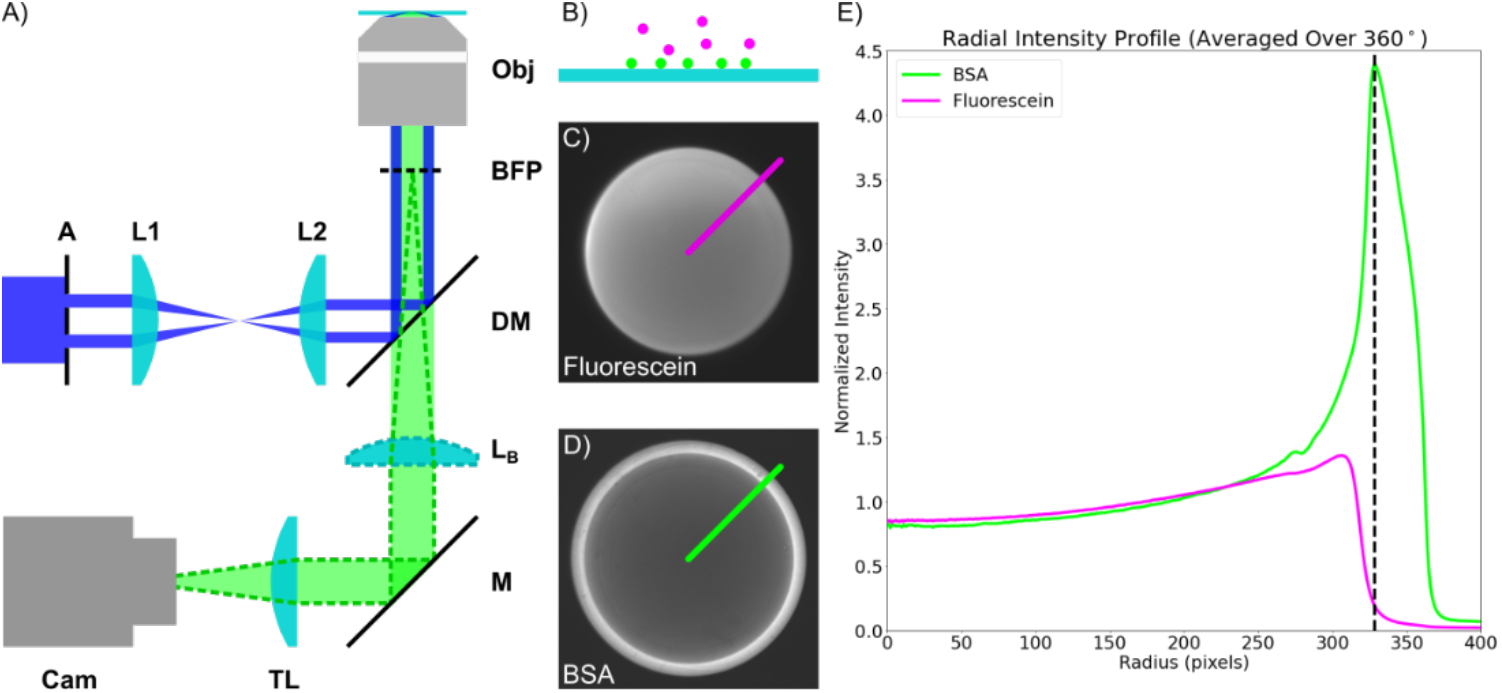
A schematic of the supercritical angle fluorescence (SAF) module. A) The sample is illuminated using a ring-TIRF configuration (blue, A) through lens pair L1 & L2. In conventional measurements, fluorescence (green) is captured by the objective and projected to the camera using a tube lens (TL). In order to capture the SAF pattern, a Bertrand lens (dotted green, L_B_) is inserted between the objective and tube lens to project the BFP of the objective onto the camera. B) Representation of fluorophores in bulk (magenta) and bound to the surface (green). C,D) Respective SAF images of fluorescein and AF-488 BSA. E) Plot of intensity for the radii indicated in C (magenta) and D (green), averaged across the images. The supercritical radius (dashed line) is found at 328 px, as identified by the intensity peak on the AF-488 BSA profile.

SAF imaging was accomplished by the addition of a Bertrand lens (AC254-200-A-ML, Thorlabs, USA) with the lens focused at the BFP of the TIRF objective. The dual deck design of the IX83 allows positioning of the Bertrand lens inside the microscope body in order to take advantage of the microscope’s tube lens and project the BFP image onto the camera. The addition of a servo motor allowed for facile switching between imaging the sample and the BFP of the objective. All images were acquired using an sCMOS camera (Prime, Photometrics, USA) and the open-source package, μ-Manager^12^.

The SAF images were processed using a Python module that locates the SAF pattern in the image and extracts the average intensity profile along the radius of the circle. The extraction process starts with a Hough’s Circle transform to identify the center of the circle, which is then cropped to the detected radius of the circle with a buffer of 50 px. An average radial fluorescence intensity was then calculated from the radial profiles about the center of the circle. Intensities at radial positions below the critical radius are considered sub-critical while those beyond the critical radius are considered super-critical. As can be seen in Figure 2, the critical radius can be readily identified from the peak in the radial fluorescence profile of surface-adsorbed species.

### C. Teflon- AF Coated Micro-Cover Glass and Characterization of AF-488 Adsorbed to Teflon-AF Film

Teflon-AF films (RI=1.35) were prepared by spin-coating Teflon-AF solutions onto 22×22 mm No.1 micro-cover glasses (~0.15 mm thick, RI=1.53) in different conditions (Table S1), after which they were heated on a hot plate at ~170°C for 10 minutes. Film thicknesses were measured by AFM using an SNL-10-B tip (Bruker, Resolve) to probe surfaces that were lightly scored to expose the glass beneath the Teflon. For most experiments, films were formed from 0.5 % w/w solutions in FC-40 at 3,000 rpm, which corresponded to ~7 nm thickness. To characterize the effect of Teflon-AF film thickness on SAF emission of AF-488, Teflon-AF coated glasses with various thickness (Table S1) were exposed to a 40 μL droplet of a 0.15 μM AF-488 BSA for three minutes and imaged with 500 milliseconds exposure time.

### D. Characterization of SAF Image Profile for Surface-Bound and Free Fluorophores

Seven standard solutions of biotin-fluorescein and free fluorescein were prepared with varying biotin-fluorescein:free fluorescein molar ratios ranging from 0 to 1, at a total fluorescein concentration of 1 μM. In each case, 40 μL of the solution mixture was pipetted onto a neutravidin-coated cover glass, followed by a 3-minute incubation period. No significant change in volume was observed during the duration of incubation and measurement. SAF data were collected by exciting the sample with 488 nm laser light and collecting images with a 100 ms exposure time. For each solution, images of five random locations on the cover glass were acquired and used to generate mean sub- and supercritical angle fluorescence profiles. Following a similar procedure, characterization of the background leakage was performed using a series of standards prepared from admixtures of 1 μM biotin-fluorescein with 0.1 to 100 μM fluorescein solution. As above, 40 μL of the admixture was pipetted onto the cover glass, followed by a 3-minute incubation period and imaging of the SAF pattern.

### E. AF-488 BSA Adsorption to Teflon-AF Coated Glass in the Presence of Surfactants

A 0.15 μM solution of AF-488 BSA, the positive adhesion control, was prepared by diluting a 76 μM stock solution in DPBS. Surfactant-BSA mixtures were prepared by mixing dilutions of stock 10 % w/w surfactant solutions (L62, L64, F68, P104, P105, F108, T904, and T90R4) with a 76 μM BSA stock solution to final surfactant concentration at 0.1 % w/w with 0.15 μM BSA. A negative-adhesion control solution, 1 μM fluorescein in DPBS, was prepared by diluting 100 μM fluorescein solution in DPBS. The negative adhesion control was used to determine the SAF profile in the absence of surface-bound fluorophores.

To measure the SAF profile, 20 μL of the sample solution was pipetted onto a Teflon-AF coated micro-cover glass (with thickness ~7 nm), followed by a 3-minute incubation time. The droplet was excited with 488 nm light and a 500-ms exposure image acquired. In each experiment, five replicate measurements were made. To determine BSA adsorption on the Teflon-AF surface, the mean SAF intensities from each sample solutions were compared to those from the negative-adhesion control sample (1 μM fluorescein in DPBS). Sample solutions with mean SAF intensities less than the negative-adhesion control were indicative of negligible BSA adsorption to the Teflon-AF layer.

### F. Monitoring of AF-488 Adsorption on Teflon-AF Coated Glass

The adsorption process of AF-488 BSA onto Teflon-AF coated glass was monitored by continuously imaging a droplet of DPBS buffer after merging it with a droplet of either (i) 0.30 μM AF-488 BSA in DPBS or (ii) 0.30 μM AF-488 and 0.1 % w/w F68 in DPBS. In brief, a 20 μL droplet of DPBS buffer was pipetted onto a Teflon-AF coated cover glass (thickness ~7 nm). The droplet was then excited with 488 nm light and images were acquired using the “multi-dimension acquisition” option in μ-Manager with the exposure time set to 10 milliseconds, interval set at 100 milliseconds, and the number of data points set at 3600 (6-minute of total acquisition time). Within the first 10 seconds of beginning the imaging, a 5 μL of sample was carefully deposited adjacent to the DPBS droplet such that the two solutions merged with minimal disturbance. This process was continuously imaged until reaching the pre-set acquisition time. All measurements were repeated three times. Both adsorption rate constant (k_a_) and half-life (t_1/2_) of AF-488 BSA on Teflon-AF were computed using pseudo-first-order equation described in the work of Alkan and coworkers^13^. Detailed calculations are included in the supplementary information.

## III. RESULTS AND DISCUSSIONS

### A. Characterization of SAF Image Profile for Surface-Bound and Free Fluorophores

To characterize the SAF imaging setup’s ability to discriminate between surface and bulk fluorescence, we acquired SAF measurements of various concentration ratios of surface-bound and free fluorophores using a model system comprised of neutravidin-coated cover glass and droplets of biotin-conjugated fluorescein and free fluorescein. The surface-bound fluorescence is controlled by the amount of biotin-conjugated fluorescein bound to the neutravidin, while the bulk fluorescence is controlled by the amount of free fluorescein in the droplet. As shown in Figure 3A, the supercritical intensity first increased with increasing molar ratio of biotin-fluorescein conjugate because of increased biotin binding to the neutravidin on the glass surface; however, the supercritical intensity plateaued when the molar ratio increased beyond 0.7. On the other hand, the subcritical intensity decreased with increasing molar ratio of biotin, which was expected as the subcritical intensity was related to the concentration of free fluorescein (bulk fluorophore). The subcritical intensity did not decrease to zero as it is likely that some of the unbound biotin-conjugate contributed to the subcritical intensity.

**Figure 3.**
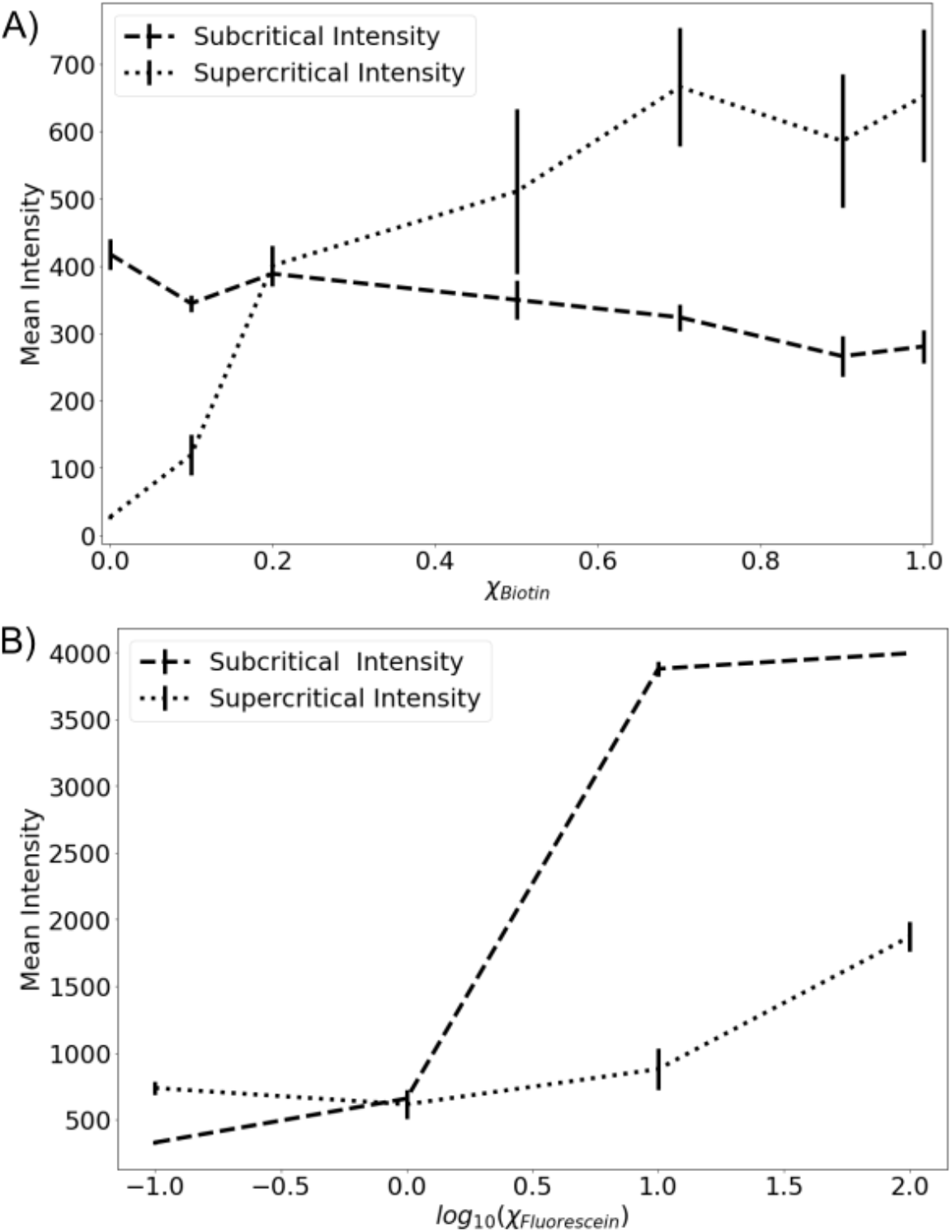
The effect of concentration of surface-bound and free fluorophores on the super- and sub-critical emission intensities. A) Plot of mean super- (short dash) and sub-critical (long dash) intensity as a function of the molar ratio of surface-bound (biotin-fluorescein conjugate) to free fluorophore (fluorescein), X_Biotin_, on a neutravidin-coated cover glass. B) Plot of mean super- (short dash) and sub-critical (long dash) intensity of solutions with constant surface-bound fluorophore concentration (biotin-fluorescein conjugate 1.0 μM) as a function of the logarithm of free fluorophore (fluorescein) concentration on a neutravidin coated cover glass. Error bars represent ± 1 std. dev. from n = 3 measurements.

We further investigated the system’s ability to differentiate between surface and bulk fluorescence. In particular, we were interested to see whether increasing the free fluorophore concentration (while maintaining constant concentration of biotinylated fluorophore) resulted in any bleed-through into the super-critical region. The changes in the supercritical and subcritical intensities as a function of total fluorescein concentration are shown in Figure 3B. The mean supercritical fluorescence intensity was first maintained at intensity 600-700 with a slight increase in subcritical intensity when the molar ratio of free fluorescein:biotin-conjugate increased from 0.1 to 1 (corresponding to 0.1 and 1 μM fluorescein). However, the supercritical intensity increased to a greater extent when molar ratios increased to 10 and 100 (corresponding to 10 and 100 μM fluorescein). This increase in supercritical intensity could be attributed to bleed-through of sub-critical fluorescence into the supercritical region. Failing to account for this effect could lead to the improper measurement of surface-adhesion events and dynamics. Albeit preliminary, our data suggests that, at least in the context of these model fluorophores, the concentration of free fluorophore should be less than 10 times that of the surface-bound fluorophore in order to minimize bleed-through effects.

### B. SAF Characterization of Adsorbed AF-488 BSA as a Function of Teflon-AF Coating Thickness

Digital microfluidics (DMF) is a liquid handling technique that manipulates microliter droplets on a Teflon-AF and dielectric coated electrode array^14^. DMF has been applied to various biology-motivated applications^14–17^, which exposed the Teflon-AF film to the protein-rich aqueous solutions. The proteins in these solutions can adsorb onto the Teflon-AF film spontaneously through non-specific adsorption (NSA), which can prevent droplet movement on the DMF platform^18^. This NSA phenomenon is challenging to study by common surface-sensitive techniques such as surface mass spectrometry ^19–21^, surface plasmon resonance^22–24^, and quartz crystal microbalance^25–27^. Many of these tools do not lend themselves readily for *in situ* measurements on substrates bearing polymer coatings, which are essential to monitor NSA on DMF. We attempted to monitor the NSA of proteins on glass substrates bearing Teflon-AF films *in situ* using SAF microscopy using a model protein, fluorescently labeled BSA, which is known to adhere to Teflon-AF. A key question in this study was the effect of film layer thickness on the SAF measurements. This is particularly important in this context as the refractive indices of Teflon-AF and BSA solution have a similar value (RI ≈ 1.3) and as such, cannot be differentiated in SAF. As illustrated in Figure 4, Teflon-AF film thickness did affect the apparent SAF intensity of adsorbed AF-488 BSA with the supercritical intensity following an exponential decay with increasing Teflon thickness. Thus, for maximum sensitivity, we elected to use the thinnest Teflon-AF layer (~7 nm) for the remainder of the experiments described here.

**Figure 4.**
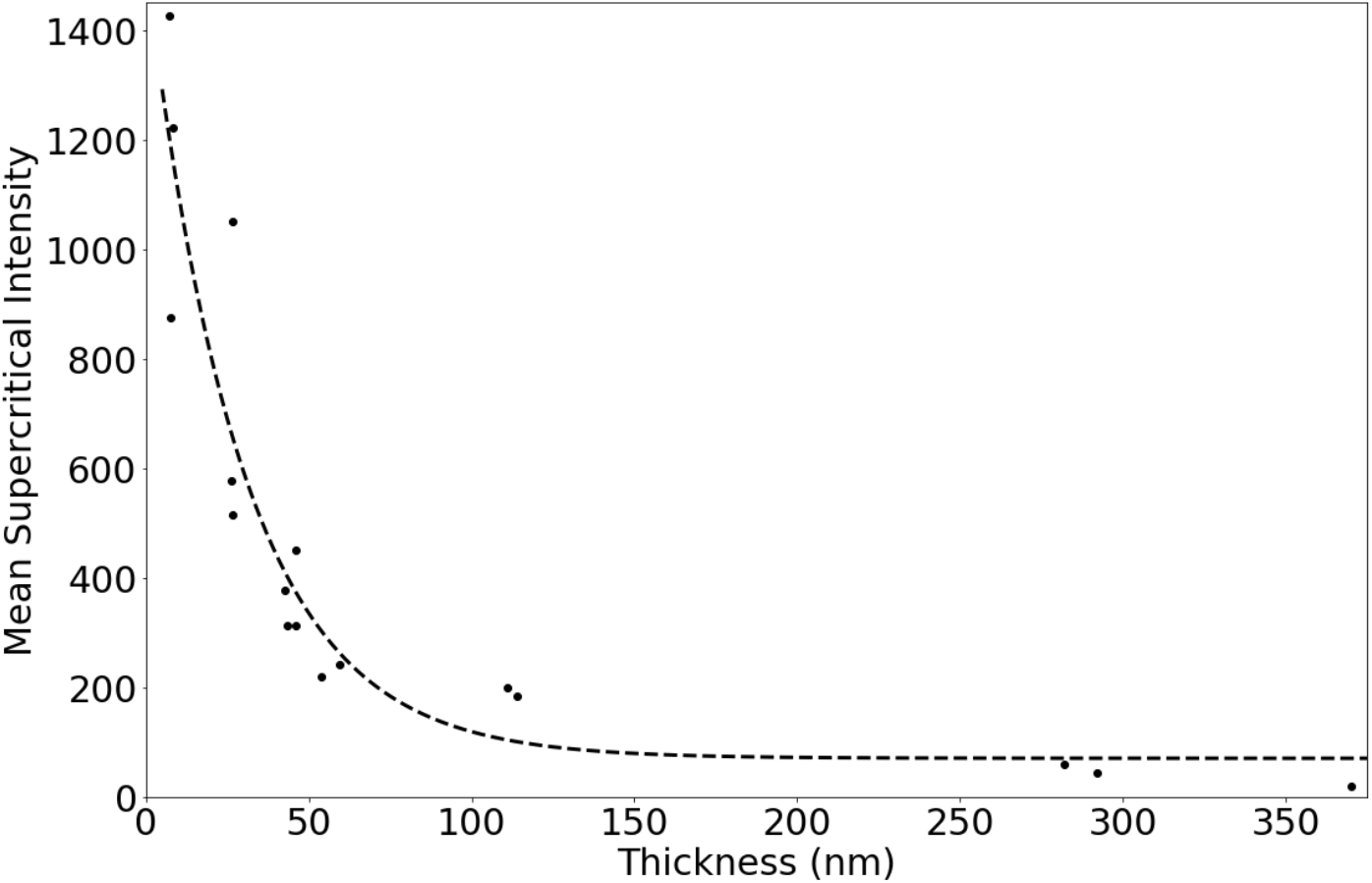
The effect of thin-film thickness on super-critical intensity. Plot of supercritical intensity (dots) of surface-bound fluorophore (fluorescently labeled BSA) as a function of film thickness of Teflon-AF (RI of 1.35). The dashed line represents a best-fit through the data

### C. Observing Non-Specific Adhesion of AF-488 BSA on a Teflon-AF Surface in the Presence of Surfactants

To combat NSA of proteins in DMF, a small quantity of Pluronic or Tetronic surfactants is commonly added to protein-rich solutions ^17, 18, 28, 29^. However, the efficacy of surfactants in combating NSA is challenging to quantify on the DMF platform *in situ* for several reasons, including (1) presence of a hydrophobic polymer thin film; (2) co-existence of liquid and solid phases, and (3) rapid interchange kinetics between surfactant, surface, and proteins. The new SAF microscopy approach described here addresses these requirements and provides a robust and straightforward technique for *in-situ* monitoring of NSA.

We analyzed the adsorption of fluorescently labeled BSA to Teflon-AF coated glass substrates in the presence of Pluronic and Tetronic surfactants to simulate the protein-surface, and protein-solution interactions typically present on a DMF platform. As shown in Figure 5, the mean supercritical intensities of each surfactant-BSA mixture did not exhibit supercritical fluorescence intensity higher than the negative-adhesion control (fluorescein), implying negligible BSA adsorption to the Teflon-AF layer. The results confirmed the previous findings of surfactants reducing BSA adsorption on the Teflon-AF layer in DMF. ^18, 28^ This is the first report of measuring surface adsorption on a thin-film polymer using SAF microscopy.

**Figure 5.**
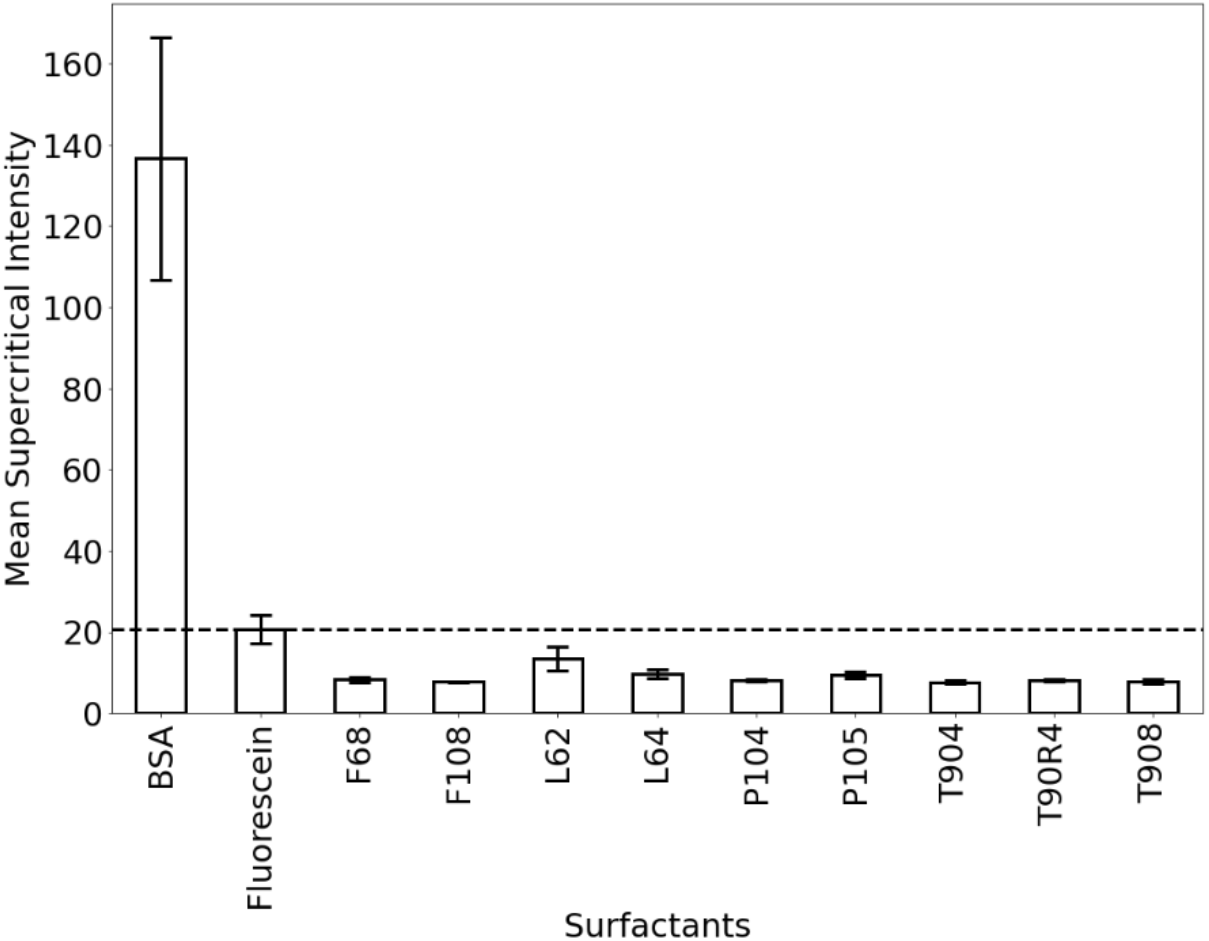
The effect of 0.1 % w/w Pluronic/Tetronic surfactants on SAF measurements on Teflon coated cover-glass. Plot of supercritical fluorescence intensity of a fluorescently labeled BSA containing no surfactant (“BSA”, left), a free fluorophore (“Fluorescein”), or labeled BSA containing surfactant (“F68” etc., right). Samples with intensity values higher than fluorescein suggested adhesion of BSA on the Teflon surface. Error bars are ±1 SD.

### D. Monitoring of AF-488 BSA Adsorption on Teflon-AF Coated Glass by SAF

Fig. 5 demonstrates a unique feature of SAF – the capacity to compare NSA *in situ* on polymer-film-coated glass substrates. This led us to hypothesize that SAF might also be able to make direct measurements of surface adsorption dynamics. With this in mind, a surface adsorption experiment was devised to evaluate the effects of exposing Teflon-AF to AF-488 BSA (Figure 6). The Teflon-AF surface was first exposed to a DPBS buffer, followed by careful merging of the original droplet with a droplet of AF-488 BSA solution. As shown, this caused an increase in the sub-critical intensity from 0 to 350 between 4-20 seconds post-merger, which we interpret to reflect the distribution of the BSA through the merged droplet. Likewise, there was an increase in super-critical emission intensity from 0 to 550 over a similar time scale, which we interpret to reflect BSA adsorption to the surface. Both super- and sub-critical intensities reached a plateau after 20 s, implying both the surface adsorption and volumetric diffusion processes reached equilibrium at that time. The gradual decrease in both super- and sub-critical intensities after 30 s can be accounted for by photobleaching. The kinetics of BSA adsorption were approximated by fitting a pseudo-first-order equation^13, 30, 31^ to the super-critical intensity data, assuming that the adsorption of BSA followed a Langmuir adsorption isotherm. The adsorption rate constant (k_a_) in our droplet merging experiment was determined to be 0.2093±0.002 s^−1^ and with half-life (t_1/2_) of adsorption 3.312±0.032 s (Fig. S1).

**Figure 6.**
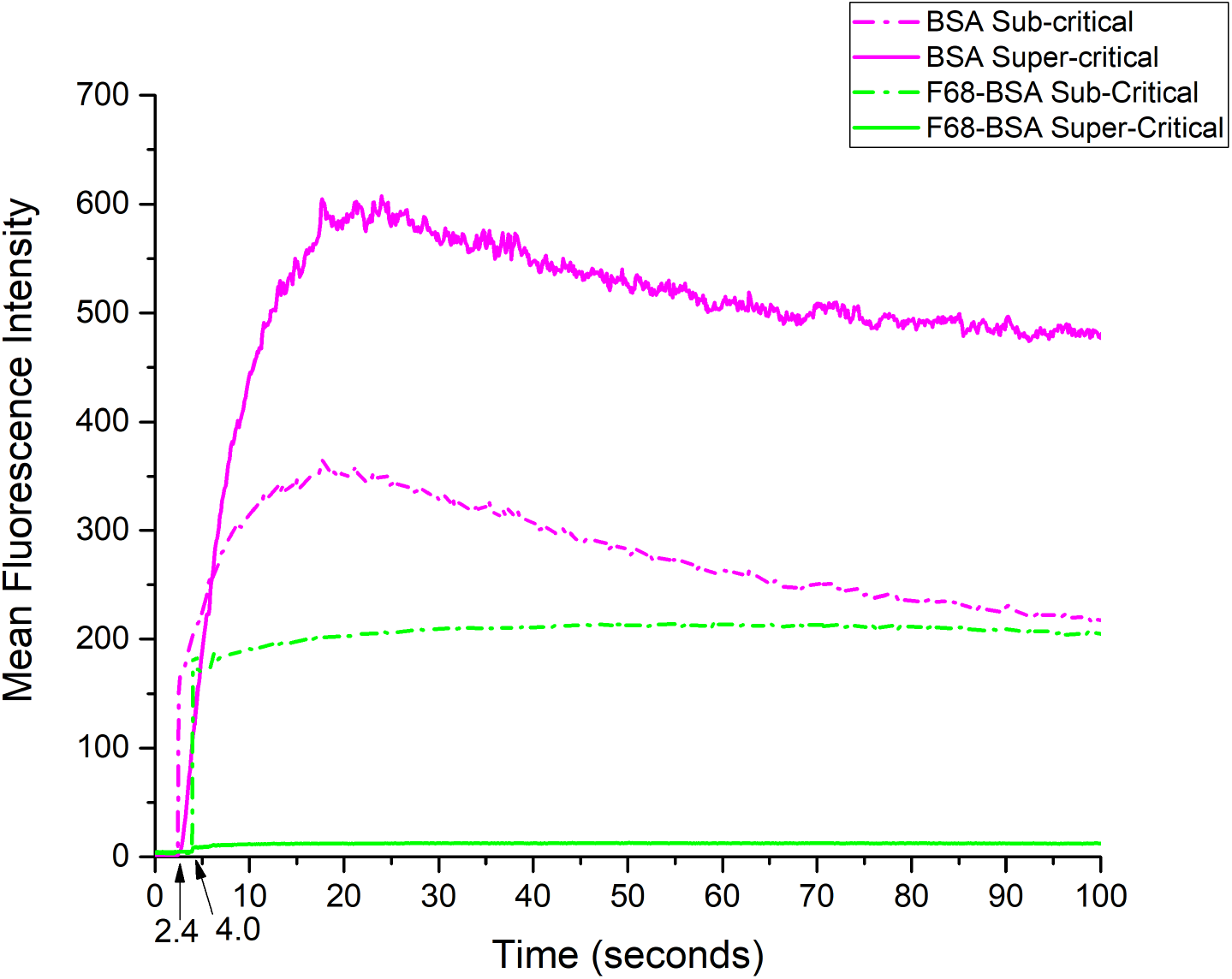
Adsorption kinetics of fluorescently labeled BSA onto Teflon-AF films. Representative plots of mean supercritical (solid lines) and sub-critical (dash-dots) intensity measured from Teflon-AF surfaces bearing a droplet of DPBS buffer after merging with a droplet of AF-488 BSA (pink) or F68/AF-488 BSA mixture (green) as a function of time. The AF-488 BSA solution was introduced at 2.4 s, while F68/AF-488BSA mixture was introduced at 4.0 s. Three measurements were done for each sample.

After verifying the kinetics of BSA adsorption without surfactant, we repeated the experiment in the presence of F68. As shown, upon merging, the sub-critical intensity increased from 0 to 200 between 5-10 seconds, which we interpret to reflect the diffusion of BSA throughout the droplet. In contrast, there was a small increase in the supercritical intensity from 0 to 10 over a similar time scale, which we interpret to mean that minimal BSA adsorbed to the Teflon-AF surface. Both super- and sub-critical intensities reached a plateau after 10s, implying volumetric BSA diffusion reached equilibrium.

## IV. CONCLUSIONS

We presented an optical module that is straightforward to build and use that enables of the observation of solid-liquid interfaces with high temporal resolution by super-critical angle fluorescence (SAF). Quantitative analysis of the SAF profiles allowed for direct determination of changes in fluorophore concentration at the surface and in the bulk solution. Control experiments that compared surface-bound and free solution fluorescence revealed that the mean SAF intensity increased with increasing concentration of surface-adhered fluorophore. However, increases in free fluorophore concentration led to sub-critical fluorescence leaking into the supercritical angle region, an effect that was only observed when the free fluorophore concentration was ≥ 10 times of surface-adhered fluorophore. In experiments with thin polymer films on glass, increasing the film thickness resulted in decreased SAF signal. Finally, the new system was used to follow the adsorption dynamics of a model protein that served as a proxy for a microfluidic device. This allowed, for the first time, the monitoring of the effects of protein in this system *in situ*, suggesting great potential for this technique for additional studies in the future.

Finally, the facile nature of the new SAF module suggests the potential for straightforward improvements in the future. For example, integration of filters or a monochromator should allow for multi-colour SAF measurements, allowing one to characterize exchange dynamics and competitive adsorption phenomena while polarized SAF measurements may provide novel insights into emission dipole orientation ^10, 32^. Likewise, the integration of a digital mirror device (DMD) in the SAF module could allow selective inspection of specific regions of interest on device surfaces, resulting in SAF measurements at specific locations of the sample. In sum, we propose that in the near future, the SAF module will be a versatile addition to any microscope permitting *in situ* monitoring of the solid-liquid interface.

## ACKNOWLEDGMENTS

This work was funded by the Natural Sciences and Engineering Research Council of Canada Discovery Grants: RGPIN-2015-043 (CMY) and RGPIN-2019-04687 (ARW).

## AUTHOR CONTRIBUTIONS

A.Au and M.H. conceived of the idea for monitoring fluorescent-labeled protein adsorption on a thin-film Teflon using supercritical fluorescence angle measurements. All reagents, micro-cover glass slides, and Teflon coatings were prepared by M.H. in the A.R.W Lab. All SAF experiments were performed by A. Au and M.H. in the C.M.Y. Lab. Atomic force microscopic measurements and SAF data analysis were all performed by A.Au. The SAF microscopic set-up was built and optimized by A.Au in the C.M.Y Lab. A. Au and M.H. interpreted results and wrote the manuscript with assistance from all authors.

## DATA AVAILABILITY

The data that support the findings of this study are available from the corresponding author upon reasonable request.

## APPENDIX: THICKNESS OF TEFLON-AF COATING AND CALCULATION OF BSA ADSORPTION KINETICS

**Table S1.** Spin coating condition and thickness of Teflon® AF layer on cover glass

| Concentration of Teflon® AF (% w/w) | Solvent | Spin speed (rpm) | Average thickness $\pm 1SD^*$ (nm) |
| --- | --- | --- | --- |
| 0.5 | FC-40 | 3000 | 7.66±0.48 |
| 1 | FC-40 | 1000 | 26.5±0.16 |
| 1 | PFC110 | 2000 | 44.0±1.43 |
| 1 | PFC110 | 1000 | 53.1±5.5 |
| 1.95 | FC-40 | 3000 | 112.5±1.5 |
| 1.95 | FC-40 | 1000 | 315±39 |

\*Thickness determined by AFM

### Calculation of adsorption rate constant (k_a_) and half-life (t_1/2_)

The integral pseudo-first-order equation is expressed as follows^13^,

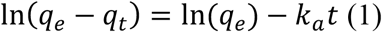

where *q_e_* and *q_t_* are the quantity of BSA (mg/g) adsorbed on a surface at equilibrium and at any time *t*, respectively, and *k_a_* is the rate constant of the pseudo-first-order-adsorption (s^−1^). The supercritical intensity is proportional to the quantity of AF-488 BSA adsorption on the surface, and we substitute *q_e_* and *q_t_* by the supercritical intensities at equilibrium and at time *t* (*I_e_* and *I_t_*) to give linearized equation (2).

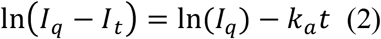

The equilibrium intensity is determined from data like that in Figure 6, where the supercritical intensity of BSA first reached plateau at ~20s. *I_q_* was then determined as the average supercritical intensity between 20-21 s. The supercritical intensity between 2.4s and 20s was computed to determine the value of ln(*I_q_-I_t_*), which was plotted as a function of adsorption time (time elapsed after addition of BSA solution) in Figure S1.

**Figure S1.**
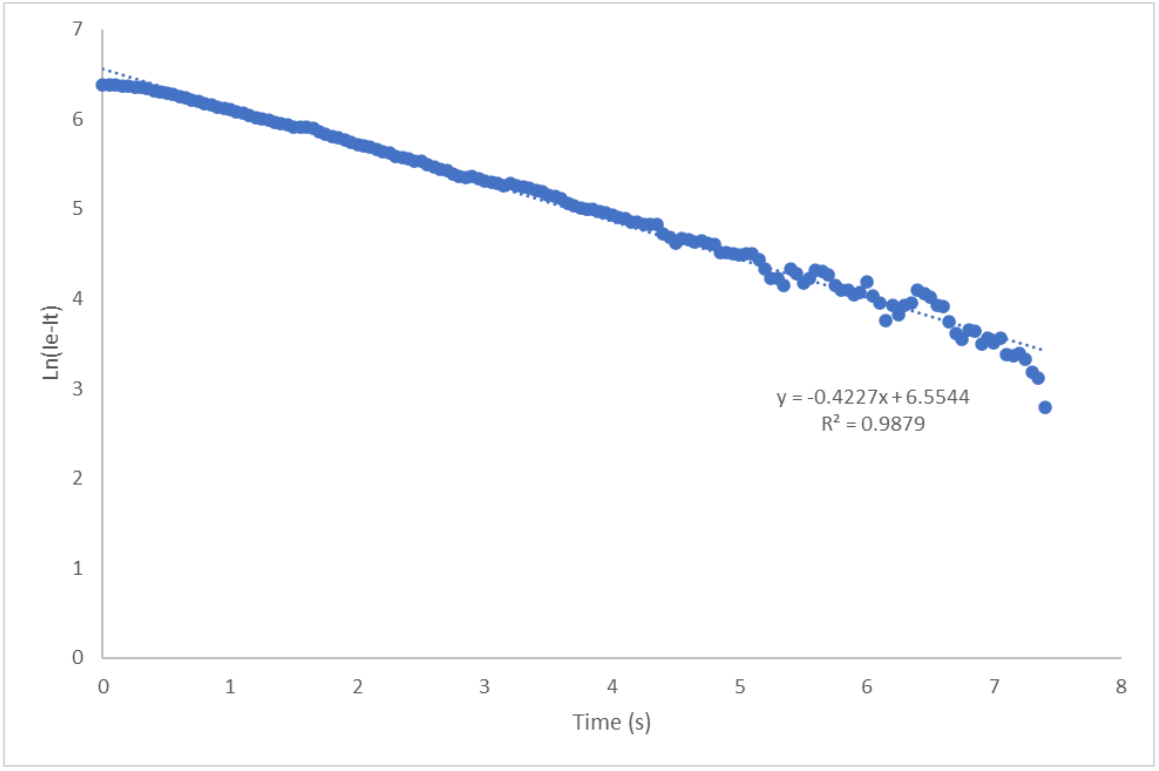
Integrated Plot for pseudo-first-order adsorption kinetic for BSA. Plot of ln(*I_q_-I_t_*) as a function of adsorption time. The adsorption rate constant (*k_a_*) was determined to be 0.4227± 0.004s^−1^ from the slope of best fit.

By equation (2),

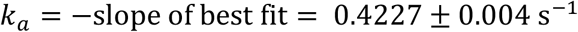

The adsorption half-life (*t_1/2_*) was calculated as,

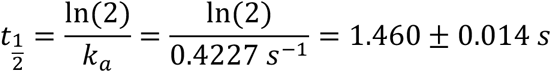

